# DECONVersation: Single-Cell Foundation Model-Derived Embeddings for Robust Cell Type Deconvolution of Bulk RNA-seq

**DOI:** 10.64898/2026.10.02.756356

**Authors:** Ali Oku, Heather Geiger, Nicolas Robine, Rui Fu

## Abstract

**Motivation:** Bulk RNA-seq deconvolution uses single-cell references to estimate cell type proportions in bulk tissue. Existing methods face challenges with batch effects, subject-level variation, and marker gene selection. We propose a new approach leveraging general-purpose single-cell foundation models.

**Results:** DECONVersation combines pretrained or fine-tuned single-cell foundation model embeddings with non-negative least squares for deconvolution. The embeddings eliminate the need for explicit marker selection, and reduce batch effects. Compared against MuSiC, DWLS, and BayesPrism, DECONVersation with Geneformer or Cell2Sentence-2B achieved competitive performance across six bulk datasets with ground truth, with performance improvements after reference-specific fine-tuning.

**Availability and Implementation:** DECONVersation is available through the Python Package Index. The source code is available at https://github.com/nygenome/DECONVersation, and the datasets and Apptainer/Singularity images are archived on https://zenodo.org/records/22949405.

**Supplementary information:** Supplementary data are available online.

## INTRODUCTION

Bulk RNA sequencing measures the genome-wide expression profile of a heterogeneous sample, from which cell type composition can be estimated through computational methods of cell type deconvolution. Deconvolution is widely used to characterize tumor microenvironments (TME) and disease-associated changes in cellular composition^1^. Single-cell RNA sequencing (scRNA-seq) has dramatically improved our understanding of cell types and their context-specific changes, but single-cell and single-nucleus isolation remain technically challenging for certain tissues^2^ and can bias the recovery of sensitive cell types^3^. In contrast, bulk profiling remains economical, easy to process, and more compatible with banked samples. Recent bulk RNA-seq deconvolution approaches attempt to bridge the two technologies, using single-cell references to infer composition from bulk data.

Deconvolution methods typically involve two steps. First, a cell type expression signature matrix is generated from a single-cell dataset or a manually defined set of marker genes. Then, a computational solver compares bulk expression with this matrix to infer cell type proportions. Most established non-deep-learning tools such as MuSiC^4^, BayesPrism^5^, and DWLS^6^ implement variations of this framework. As such, deconvolution performance is strongly influenced by the transferability of the signature matrix across samples and technologies. Manually curated signatures are labor-intensive and susceptible to curator bias, while scRNA-seq-derived signatures can be skewed by clustering and annotation decisions, batch effects, and donor-level variation. Existing methods attempt to address these issues differently. MuSiC weights genes by subject-level variance, BayesPrism marginalizes expression differences between reference and query in a Bayesian framework, and DWLS uses dampened weighted least squares to prioritize marker genes. However, benchmarking studies still demonstrate vulnerability to batch effects, often leading to large errors in estimated proportions^7^, and instability across marker gene selection methods^8^, for which no single marker strategy suits all tissues tested.

Here, we propose DECONVersation, which uses single-cell foundation models (scFMs) trained on large-scale single-cell data to generate embeddings that capture the context of an entire cell/sample by aggregating information across thousands of gene tokens. It currently incorporates embedding extraction from four scFMs: Geneformer^9^, Cell2Sentence (C2S)^10^, scGPT^11^, and CellHermes^12^. Because scFMs are trained on large and diverse single-cell data, their embeddings are robust to technical variation and batch effects^13^ while reflecting meaningful biology, as demonstrated by their capabilities in batch integration and cell type annotation. scFMs can learn gene networks^9–12^ directly from single-cell data, capturing biological signals while reducing noise^14^. Fine-tuning on reference single-cell data can further add tissue-specific learning to scFM embeddings, in lieu of the marker selection step of traditional tools.

We evaluated embeddings from several widely used and new prototype scFMs. Geneformer, a BERT-like transformer encoder, was pretrained on over 104 million human single-cell transcriptomes with a self-supervised masked language modelling objective, where genes within each cell are ranked by normalized expression level and the model learns to predict masked gene ranks. C2S represents genes as natural language sentences and is evaluated in two variants, C2S-410M, built on EleutherAI’s Pythia architecture, and the subsequent C2S-2B, built on Google’s Gemma 2 framework. CellHermes extends this multimodal approach, framing transcriptomic profiles, protein-protein interaction networks and phenotypic metadata as text-based question-answer pairs, and uses a Low-Rank Adaptation (LoRA)^15^, fine-tuned large language model (LLM) core to align cross-modal biological entities. In contrast, scGPT couples a specialized gene vocabulary with binned continuous expression values rather than directly ranking genes or translating expression to text, using a custom generative decoder-based transformer architecture pretrained on over 33 million cells^11^.

Although designed for and trained on single-cell data, scFMs have occasionally been explored for bulk RNA sample tasks such as marker selection^16,17^ and tissue classification^10,18^. Here, we present DECONVersation, a novel application of scFM embeddings to bulk profiles to address persistent deconvolution challenges, including selecting informative marker genes and minimizing technical and batch effects across platforms and tissue contexts. We demonstrate its efficacy by benchmarking against several existing high performance tools.

## METHODS

### Data Preparation

As all four foundation models accept single-cell input, bulk RNA-seq counts or TPM, as provided by the publication sources, were converted to AnnData format, treating samples as pseudo-cells. Foundation models were then used to generate embeddings from each pseudo-cell. Similarly, cell type signatures, average expression within cell types, and pseudobulks were derived from single-cell data, converted to AnnData format, and then used for embedding extraction. Geneformer required Ensembl IDs for gene representation, while CellHermes, scGPT and C2S used gene symbols.

### scFM Embedding Extraction and Fine-tuning

All embedding extraction was done in Python (v3.10), PyTorch^19^ (v2.1) and HuggingFace Transformers (v4.36), with parameters not stated below kept at the authors’ recommended defaults.

For Geneformer (v2), tokenization used TranscriptomeTokenizer with a 4,096-token input size, special tokens enabled, and a chunk size of 512. Cell-level embeddings were extracted with get_embs (emb_mode=“cell”) from the final layer of the 18-layer V2 architecture, with a forward batch size of 50 to accommodate GPU memory constraints. We modified the code to support CPU and Apple Silicon (MPS) extraction.

C2S (PyPI v1.2.0) was used for the 410M (C2S-410M) and Scale 2B (C2S-2B) models. Transcriptomic data were converted to cell sentences with the adata_to_arrow function, and embeddings were extracted with embed_cells, modified to support MPS, using 200 genes per cell sentence. We added LoRA fine-tuning to enable fine-tuning of 2B models on memory-constrained GPUs.

The CellHermes V1 model was loaded in inference mode using a Low-Rank Adaptation (LoRA) configuration with the LLaMA-3 prompt template. Following the authors’ examples, mitochondrial and ribosomal genes were excluded, and the top 500 ranked genes per cell were formatted as a structured natural language prompt. The final hidden state of the last token was used as a 4,096-dimensional cell embedding. Batch embedding and fine-tuning functions were adapted from the CellHermes example scripts.

scGPT (v0.2.5) embeddings were extracted with the tasks.embed_data function, with gene identifiers supplied via gene_col and a batch size of 64. Because bulk profiles lack the sparsity of single-cell data, we modified the code to tokenize the top 1,200 expressed genes rather than randomly downsample.

All models were fine-tuned to predict annotated cell types from the reference, following each model’s examples and utility code. As several scFMs have been shown to suffer in performance when fine-tuned on imbalanced data, over-represented cell types were randomly undersampled before fine-tuning^20^.

For comparison, scVI embeddings were extracted using scvi-human-2024-07-01^21^ which was pretrained on human single-cell data from the CZ CELLxGENE Census. Bulk RNA-seq data and pseudobulked cell type signatures were aligned to the model’s reference genes and loaded as query data. A forward pass generated 50-dimensional sample embeddings without query-specific training. Additionally, we extracted principal component scores for each signature matrix and its corresponding bulk expression matrix by fitting PCA to the signature matrix using scikit-learn^22^ and projecting the bulk matrix onto the resulting components.

### Bulk RNA-Seq Benchmarking

We benchmarked DECONVersation across six bulk RNA-seq datasets with ground truth proportions from previous benchmarking studies, including three peripheral blood mononuclear cell (PBMC) cohorts^23–25^, a retinal tissue dataset^26^, a pre-frontal cortex dataset^27^ and a cell line mixture dataset^28^. The PBMC cohort included paired bulk RNA-seq and ground-truth fractions from fluorescence-activated cell sorting (FACS). Ground truths were derived from known mixing counts for the cell line mixture, and estimated from sample-matched snRNA-seq and RNAscope for the retinal and pre-frontal datasets, respectively.

Pre-processed single-cell references were obtained from the same studies as their corresponding bulk datasets^29,30^. For the cell line mixture, clear doublet clusters observed during visualization were removed with scDblFinder^31^. Except for the pre-frontal cortex data where all cells were used, cells were downsampled to the same number per cell type in each reference, with the exception of very low count cell types, ranging from 800 to 5000 depending on the study, to provide a meaningful spread of numbers representative of real-world single-cell/nuclei datasets used for deconvolution.

### Pseudobulk Benchmarking

To benchmark deconvolution against known proportions in additional contexts, pseudobulk samples were generated by randomly sampling cells from single-cell datasets and summing their raw counts. In each sample, a randomly selected target cell type was assigned a proportion drawn uniformly from 10–80%, with remaining cells drawn without constraint from the pooled non-target cell types. We generated 250 samples of 300 cells for broad cell type analyses and 500 samples of 1,000 cells for finer cell type analyses.

To estimate the effects of technical and batch differences, we varied the differences between each pseudobulk source and the deconvolution reference. For each pair, signature matrices were restricted to shared genes, the Pearson correlation was computed for each matched cell type, and dissimilarity was defined as 1 − the minimum correlation, a conservative score driven by the least concordant cell type. For endometrium, single-donor pseudobulks were generated with increasing technical dissimilarity to the reference: a different donor from the same cohort and assay (snRNA-seq), a different cohort with the same assay (snRNA-seq), and a different cohort and assay (scRNA-seq), the latter two from the Human Endometrial Cell Atlas (HECA). For PBMC, pseudobulks were generated from three public 10x Genomics datasets (5′ single-cell, single-nucleus, and probe-based Flex) and deconvolved using the PBMC reference mentioned above for real bulk data.

### Attention Extraction and Cell Type Enrichment Analysis

Gene-level attention scores were extracted from pretrained and fine-tuned Geneformer models using HuggingFace Transformers. For each cell, final-layer attention from a single forward pass was averaged across heads and the CLS token row was taken as the attention allocated to each gene. Scores were then averaged across cells to give a ranked attention profile per cell type. The top 120 genes by mean attention were used for cell-type enrichment analysis (over-representation analysis) with GSEApy^32^ and the PanglaoDB (2021)^33^ cell-type marker reference, following a similar approach to Xiao et al^34^. We used 50 cells per cell type for this analysis, though DECONVersation supports larger cell counts and broader gene rankings.

### Deconvolution Solvers

DECONVersation uses Non-Negative Least Squares (NNLS, from SciPy) by default, and supports four other solvers: Nu Support Vector Regression (nuSVR), Ridge regression (scikit-learn), gradient descent constrained cosine solver, and a Python reimplementation of dampened weighted least squares (DWLS).

### Evaluation

For each dataset, performance was evaluated by root mean square error (RMSE) and Pearson correlation, overall (Cor) and for each cell type (typeCor), between estimated and ground-truth cell type proportions. DECONVersation was benchmarked against three established methods which include MuSiC, BayesPrism, and DWLS, run with author-suggested defaults. We also computed the condition number (numpy.linalg.cond) of each reference’s raw signature, DWLS marker, and Geneformer and C2S-2B embedding matrices. The condition number measures how sensitive estimated proportions are to small changes or noise in the signature matrix, indicating how easily cell types can be distinguished.

### Software implementation

DECONVersation code and tutorials are available at https://github.com/nygenome/DECONVersation and can be installed from PyPI at https://pypi.org/project/DECONVersation/. Because the four scFMs are not compatible in the same Python environment due to dependencies, DECONVersation detects and loads only the available model(s). For each scFM and software package, users should consult the corresponding official installation guides. For reproducibility, Conda environment YAML files and prebuilt Singularity images with each scFM and DECONVersation v0.1.0 are provided at https://zenodo.org/records/22949405. GPU runtime benchmarking was performed with NVIDIA L40S.

## RESULTS

### Deconvolution performance on real bulk RNA-seq data

DECONVersation generates embeddings from bulk expression and single-cell signature matrices, then passes both directly into a deconvolution solver (**Fig 1A**). Embeddings come from either pretrained scFMs or models fine-tuned for cell type classification using the reference single-cell/nuclei (sc/sn) data. In contrast, standard tools operate in gene expression space for both the signature matrix and the bulk matrix. To demonstrate that embeddings can represent linear cell proportion relationships, we built pseudobulks from 4 cell types in 10x Genomics example 3’ PBMC data and deconvolved them against the same reference using zero-shot Geneformer embeddings **(Fig 1B)**. In this near-ideal scenario, predicted proportions were highly correlated with ground truth proportions (r > 0.98 across all cell types).

**Figure 1:**
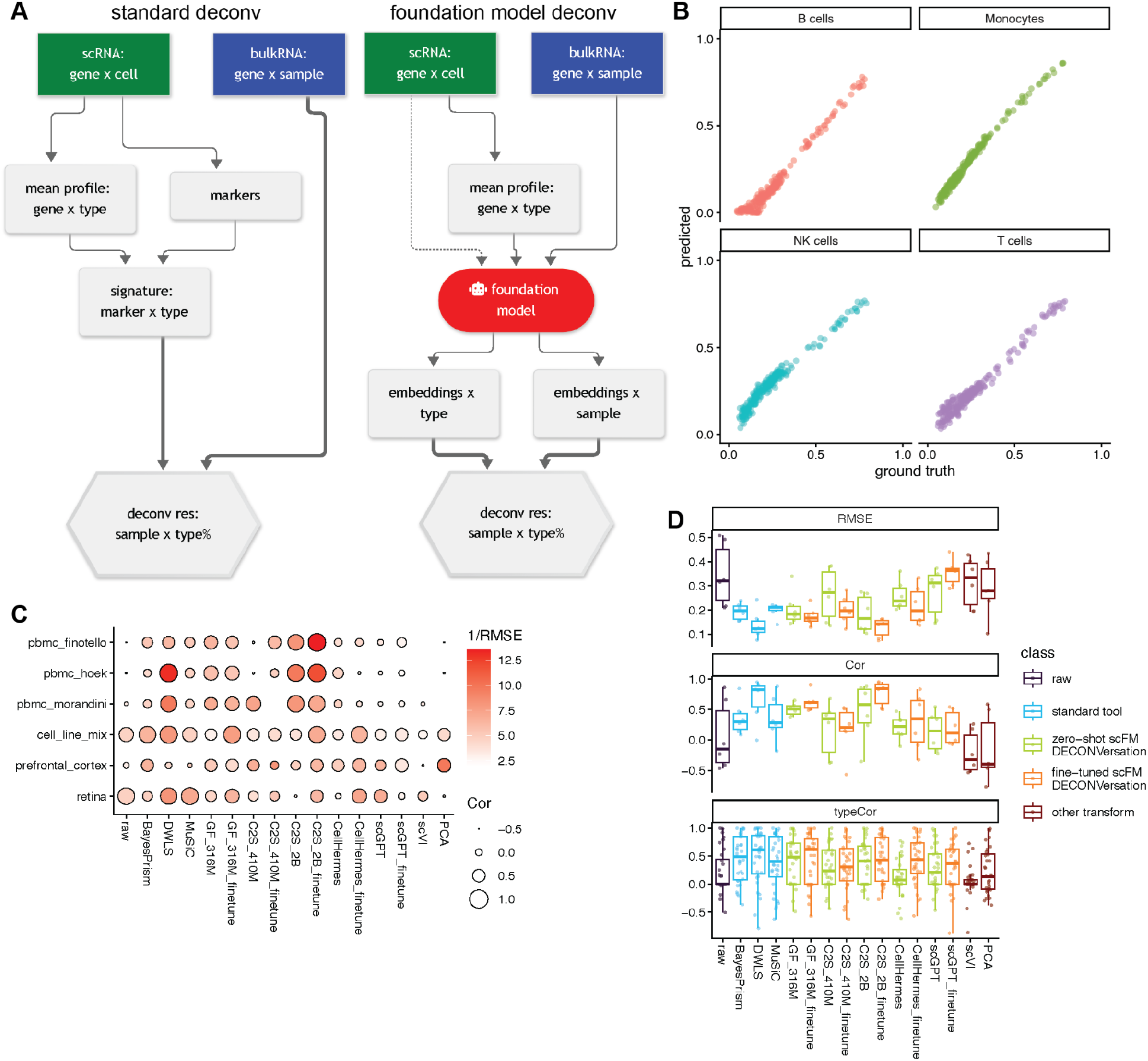
Deconvolution performance across bulk datasets. A) Workflow diagrams of conventional bulk RNA-seq deconvolution using single-cell or nuclei reference, and using scFM zero-shot or fine-tuned (dotted line) for deconvolution in DECONVersation. B) Example results from PBMC pseudobulk of four cell types. C) Dotplot of results from all 6 bulk datasets with ground truth, showing 1/RMSE (darker is better) and overall correlation (larger is better). D) RMSE and correlation, overall and per cell-type, summarized to boxplots. **ALT TEXT**: Workflow diagrams and graphs labelled A to D showing deconvolution performance. A is a workflow diagram comparing standard bulk RNA-seq deconvolution with DECONVersation. B is a scatter plot comparing ground-truth cell-type proportions against predictions for PBMC pseudobulk. C is a dot plot of six datasets displaying correlation by dot size and 1/RMSE by color intensity. D is boxplots summarizing performance metrics including RMSE, sample correlation, and cell-type correlation across tools.

We examined the performance of the scFM-based methods on real bulk RNA-seq datasets. From recent benchmarking studies, six bulk RNA-seq datasets with known composition were assembled (Table 1), and three well-performing deconvolution tools^7,35^, MuSiC (weighted NNLS), DWLS (dampened weights to address rare cell types and low-expression markers) and BayesPrism (Bayesian joint estimation of cell-type fractions and cell-type-specific expression) were selected as comparison targets. These benchmark studies also provide broader context on the performance of additional deconvolution methods. We then computed correlation, overall and per cell-type, and RMSE against ground truth, as done in previous benchmarks^7,36^. For example, in the Hoek et al. bulk PBMC dataset (Supplemental Fig S1), fine-tuned C2S-2B matched DWLS as the best-performing tool (r 0.952 vs 0.948, RMSE 0.084 vs 0.076), while all other Geneformer and C2S models outperformed BayesPrism and MuSiC. Across all bulk datasets, zero-shot Geneformer and C2S-2B (mean r = 0.50 and 0.46) outperformed MuSiC (0.35) and BayesPrism (0.36), though both trailed DWLS (0.65), the best-performing standard tool. After fine-tuning, Geneformer (0.62) approached DWLS and C2S-2B outperformed it (0.76). Standard tool performance varied by dataset. On prefrontal cortex, DWLS and MuSiC showed low average correlation (r = −0.004 and −0.181 respectively) while BayesPrism performed poorly on the PBMC datasets (Finotello: r = 0.330; Hoek: r = 0.089; Morandini: r = 0.142). Consistent with previous benchmarks, DWLS was generally the strongest standard tool. scGPT and CellHermes varied across datasets but generally outperformed raw expression coupled with NNLS (mean r 0.13 and 0.20 vs 0.06). Unlike C2S and Geneformer, their inconsistency was not resolved by fine-tuning. Notably, no tool achieved top or near-top accuracy across all datasets (**Fig 1C,D)**, highlighting unresolved issues, potentially tissue-specific constraints on marker signal-to-background strength and/or batch-driven effects.

**Table 1.** Bulk and pseudobulk datasets and references used.

| Source | Real bulk | Ground Truth | Tissue | Cell Type # | Cell Types | Bulk sample # | Count or TPM | Reference | Ref sc or sn | Reference cell # |
|---|---|---|---|---|---|---|---|---|---|---|
| Cobos et al. 2023 | Yes | mixture count | Cell line mixture | 6 | BT474,hMSC,Jurkat,MCF7,T47D,THP1 | 36 | Counts | Cobos et al. 2023. Cleaned | sc | 4140 |
| Hoek et al. 2015 | Yes | FACS | PBMC | 5 | B cells,DC,Monocytes,NK cells,T cells | 8 | TPM | Hao et al. 2021. Subset | sc | 4800 |
| Finotello et al. 2019 | Yes | FACS | PBMC | 5 | B cells,DC,Monocytes,NK cells,T cells | 9 | TPM |  |  |  |
| Morandini et al. 2024 | Yes | FACS | PBMC | 5 | B cells,DC,Monocytes,NK cells,T cells | 159 | TPM |  |  |  |
| Huuki-Myers et al. 2025 | Yes | RNAScope/IF | DLPFC | 6 | Astro,EndoMural,Excit,Inhib,Micro,OligoOPC | 81 | Counts | Huuki-Myers et al. 2024. Matching sn | sn | 56447 |
| Guo et al. 2025 | Yes | snRNA | Retina | 6 | AC,BC,Cone,HC,NN,Rod | 20 | Counts | Liang et al. 2023. From raw counts | sn | 30000 |
| Marečková et al. 2024 (sc) | No | scRNA pseudobulk | Endometrium | 9 | dStromal_early,dStromal_mid,eHormones,eStromal,Glandular,Immune_Lymphoid,preGlandular,SOX9_functionalis_II,Venous | 500 | Counts | Marečková et al. 2024 (sn) | sn | 43394 |
| Guimarães et al. 2024 (inDrop) | No | scRNA pseudobulk | Lung tumor | 10 | B cell,endothelial cell,epithelial cell,fibroblast,malignant cell,mast cell,mononuclear phagocyte,NK,TCD4,TCD8 | 500 | Counts | Guimarães et al. 2024 (10x3'v2, sc) | sc | 9563 |

Thus, embeddings from scFMs, especially Geneformer and C2S, have the potential to resolve cell type composition, on par with popular tools in the field. Fine-tuned C2S-2B demonstrated the best accuracy, while zero-shot Geneformer was also well positioned, with the benefit of forgoing fine-tuning entirely. For technical detail discussions below, we focus on these two scFMs.

### Deconvolution performance on pseudobulk data

As previous studies have repeatedly discussed, pseudobulks sampled from and tested on the same reference do not reflect real-world batch effects and technical differences between bulk and single-cell/nuclei platforms^37^. We therefore assessed assay-driven batch effects by deconvolving endometrial lineage-level pseudobulks from three sources of progressively greater technical dissimilarity against the same reference. Although performance decreased with dissimilarity for all tools, scFM-based methods generally maintained higher correlation and comparable RMSE relative to MuSiC and BayesPrism, while fine-tuned Geneformer and DWLS were the best at the highest dissimilarity (**Fig 2A**). Similarly, using a 3′ scRNA-seq PBMC reference to deconvolve pseudobulks from 5′ scRNA-seq, snRNA-seq, and probe-based Flex assays, Geneformer and C2S-2B performed on par with or better than MuSiC and BayesPrism in each scenario (**Supplemental Fig S2**).

**Figure 2.**
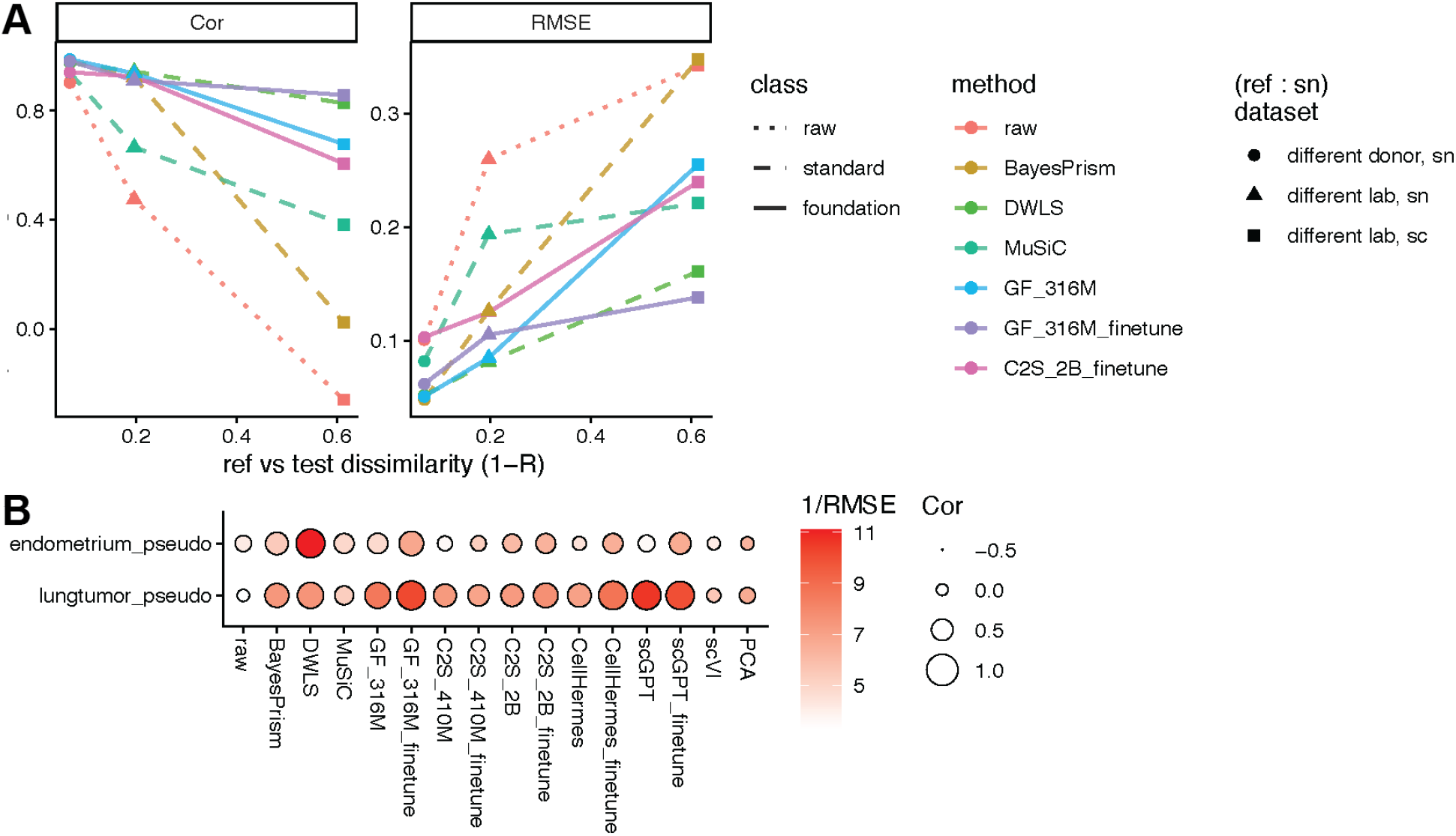
Pseudobulk benchmarks. A) Comparison of various deconvolution tools using the same endometrium reference against increasingly dissimilar single-cell/nuclei datasets, pseudobulked at lineage level. B) Comparison of various deconvolution approaches against HECA and lung TME finer cell type pseudobulk data. **ALT TEXT**: Performance graphs labelled A to B comparing deconvolution tools on pseudobulk datasets. A is a line graph comparing performance under increasing query-reference dissimilarity. B is a dot plot comparing performance on finer cell types in HECA and lung tumor microenvironment data.

Bulk RNA-seq data with real ground truths are scarce and limited by the resolution of orthogonal assays (no more than 6 broad cell types in our test datasets). To evaluate finer resolution, we deconvolved pseudobulks from a Human Endometrial Cell Atlas (HECA) scRNA-seq cohort^38^, using an snRNA-seq reference of 9 cell types across 4 major lineages. As expected, performance decreased for all methods relative to lineage level. However, fine-tuned scFMs remained competitive with established tools, underscoring the value of fine-tuning for finer-cell type use cases. Notably, fine-tuned Geneformer (r = 0.65) outperformed both BayesPrism (r = 0.55) and MuSiC (r = 0.41), while zero-shot Geneformer (r = 0.44) remained comparable to these standard tools. DWLS achieved the highest overall correlation (r = 0.86) (**Fig 2B**). We observed similar results in a 10 cell-type experiment using the lung tumor subset of an integrated pan-cancer TME atlas^39^, with inDrop data for pseudobulking and 10x 3′ v2 data as the reference. Here, fine-tuned Geneformer (r = 0.87) outperformed DWLS (r = 0.73), fine-tuned C2S-2B (r = 0.65), zero-shot C2S-2B (r = 0.53), BayesPrism (r = 0.62), and MuSiC (r = 0.36).

### Technical details

As transformer-based scFMs have many presumed strengths but limited interpretability, we explored several facets of how they suit deconvolution. First, we embedded GTEx^40^ bulk profiles with zero-shot Geneformer and C2S-2B models and projected embeddings directly on UMAP. Samples of the same tissue origins projected into nearby UMAP space, in similar fashion to UMAP projection from PCA after highly variable gene selection, showing that meaningful biological differences among bulk samples are preserved in zero-shot embeddings **(Fig 3A)**.

**Figure 3.**
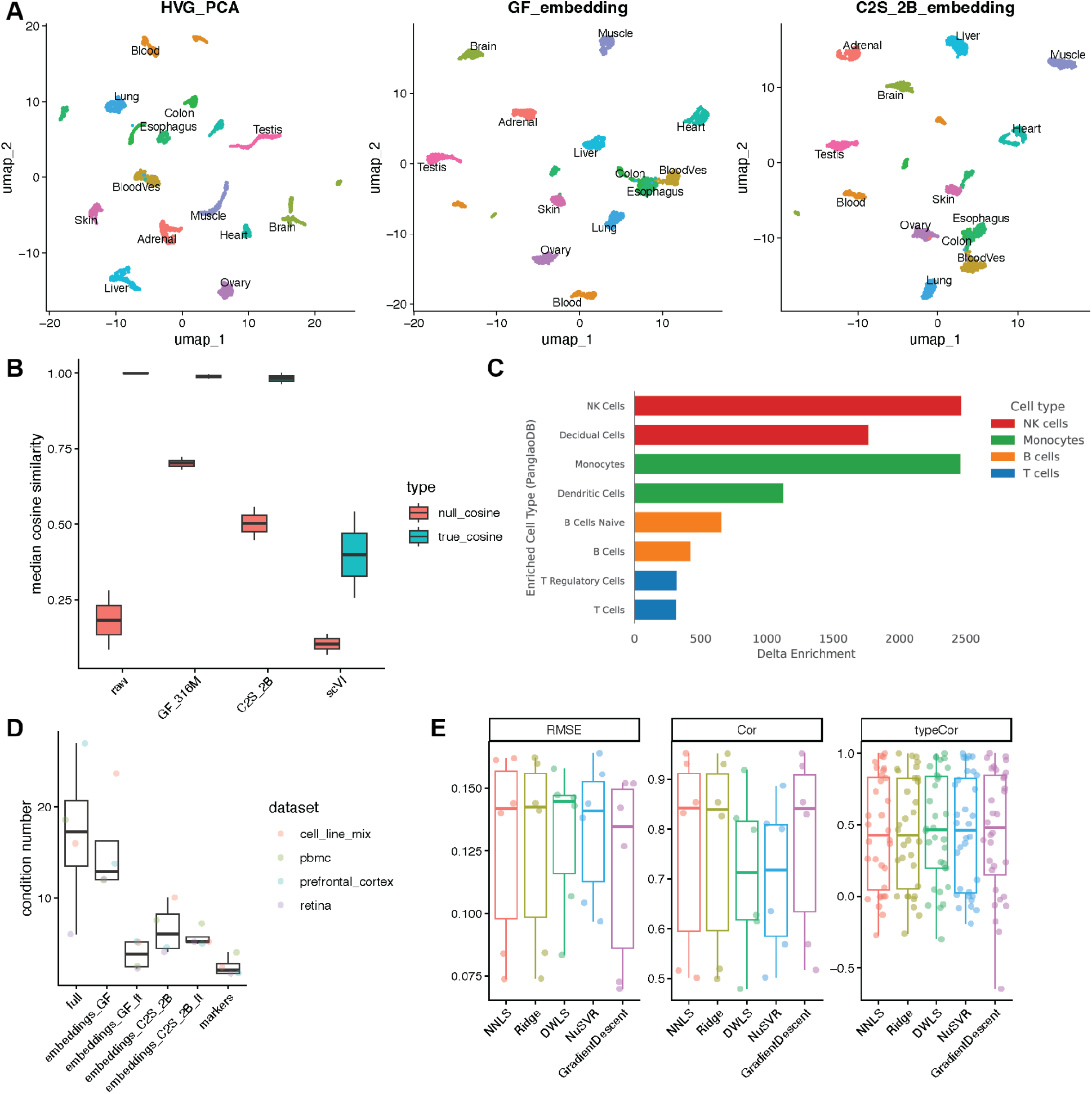
Exploration of the feasibility of scFM embeddings for bulk RNA-seq deconvolution. A) GTEx bulk RNA UMAP projections based on (left) highly variable feature selection and PCA, (middle) Geneformer 316M zero-shot embeddings, and (right) C2S-2B zero-shot embeddings. B) Median cosine similarity comparing endometrium lineage and PBMC pseudobulk embeddings to additive signature embedding sums based on ground truth proportions, or random GTEx bulk RNA embeddings as null comparison. C) Comparison of PanglaoDB marker gene set enrichment scores derived from attention-based gene extraction in zero-shot and fine-tuned Geneformer models. Delta enrichment reflects the difference in combined score between shared cell-type categories across the two models (fine-tuned minus zero-shot). D) Condition number comparison between raw data, various scFM embeddings, and DWLS-selected markers. E) Comparison of NNLS and other solvers on C2S-2B fine-tuned embeddings. **ALT TEXT**: Graphs labelled A to E. A are UMAP projections of GTEx bulk data comparing PCA, Geneformer, and C2S embeddings. B is boxplots of median cosine similarity to additive signature sums. C is bar plots of PanglaoDB marker gene enrichment score differences between zero-shot and fine-tuned models. D is a graph of condition numbers across embeddings. E is box plots comparing solver performance on C2S-2B embeddings.

Next, we assessed whether embeddings respect the linear structure expected by most solvers. Using endometrium lineage and PBMC pseudobulks, we calculated cosine similarity between embeddings generated from pseudobulk profiles and linear combinations of signature embeddings weighted by ground-truth proportions. Randomly selected GTEx sample embeddings were used as a null background (**Fig 3B**) because transformer embeddings may contain a large shared offset or dominant mean direction (anisotropy)^41^ that can inflate similarity. As expected, raw counts without transformation are naturally linear. In contrast, scVI embeddings show little anisotropy but do not combine linearly. Though strong anisotropy was observed through median cosine similarities above 0.5 on null tests in Geneformer and C2S-2B architectures, correctly matched proportions still had a meaningful linear effect compared to the null, with cosine similarities all above 0.95.

Third, to confirm that Geneformer has learned relevant biological signals, which can be further reinforced through fine-tuning, we extracted average gene attention during forward passes of 50 cells per PBMC cell type. For B cells, for example, many of the top genes ranked by attention in both zero-shot and fine-tuned Geneformer are common B cell markers (**Table 2**). Enrichment analysis on attention rankings from the fine-tuned model showed higher enrichment of corresponding or similar PanglaoDB cell type marker sets than zero-shot (**Fig 3C**).

**Table 2:** Top 10 genes ranked by model attention across cell types (zero-shot vs. fine-tuned)

| **Cell type** | **Zero-shot** | **Fine-tuned** |
|---|---|---|
| B cells | GNAS, FTL, CD79A, LAPTM5, LTB, CD52, CYBA, TXNIP, CXCR4, KLF2 | LTB, CD79A, MEF2C, GNAS, KLF6, HLA-DPB1, CD52, RPS11, TXNIP, TOMM7 |
| Monocytes | LYZ, FTL, CCL4, CTSS, CXCL8, CST3, S100A9, CYBA, LAPTM5, IL1B | CTSS, AHNAK, CD14, NFKBIA, TYROBP, PSAP, MNDA, CST3, LYZ, SH3BGRL3 |
| NK cells | GNAS, CTSW, FTL, CCL4, PTPRC, GZMB, CYBA, CALM1, GNLY, CCL5 | GNLY, CALM1, HCST, KLRF1, GZMB, PTPRC, ID2, CD7, GZMK, CTSW |
| T cells | FTL, GNAS, ZFP36L2, LTB, IL7R, PTPRC, TXNIP, CALM1, CD52, TMSB4X | LTB, KLF6, CALM1, IL7R, CD52, GNAS, KLF2, GZMK, RPLP0, ZFP36L2 |

Next, to examine whether scFM embeddings were associated with easier deconvolution solving, we calculated condition numbers, often used for marker selection optimization (including in DWLS), for the four references used in real bulk deconvolution. All embedding-based methods and DWLS had lower average condition numbers than the full gene matrix. As expected, fine-tuned scFM embeddings had a lower average condition number than zero-shot embeddings. DWLS markers were the lowest, which may partially explain DWLS’s strong performance but is also a product of its explicit marker selection optimization for the lowest condition number (**Fig 3D**). We then evaluated whether solver choice affected performance across all six bulk datasets, coupling C2S-2B fine-tuned embeddings with the five solvers in DECONVersation: NNLS, nuSVR (the CIBERSORTx regression solver), Python-reimplemented dampened weighted least squares (solver-only portion of the DWLS R package), ridge regression, and a gradient descent constrained cosine solver. Performance was generally consistent and not substantially improved by solvers with different assumptions and fitting objectives (also observed on Geneformer zero-shot embeddings, **Supplemental Fig S3**). NNLS, a fast and commonly used baseline with minimal hyperparameters, was therefore chosen as the default (**Fig 3E**).

We also evaluated which Geneformer layer provides the most appropriate embeddings for deconvolution. The Geneformer API exposes the last two layers for extraction and recommends the second-to-last by default, on the basis that the last layer is weighted toward the direct learning objective while the second-to-last is more generalizable. Our use case is different, because Geneformer learns to differentiate between cell types in both pretraining and fine-tuning, so the last layer may be more suitable. We therefore extracted embeddings from all 18 layers and tested them on the six bulk datasets. For both zero-shot and fine-tuned models, the final two layers (17 and 18) consistently yielded the best results. Layer 18 slightly outperformed layer 17, with higher average correlation and lower RMSE across the bulk datasets (**Fig 4A**). DECONVersation therefore extracts embeddings from the last layer by default, though users can choose other layers. C2S also outputs last-layer embeddings by default.

**Figure 4.**
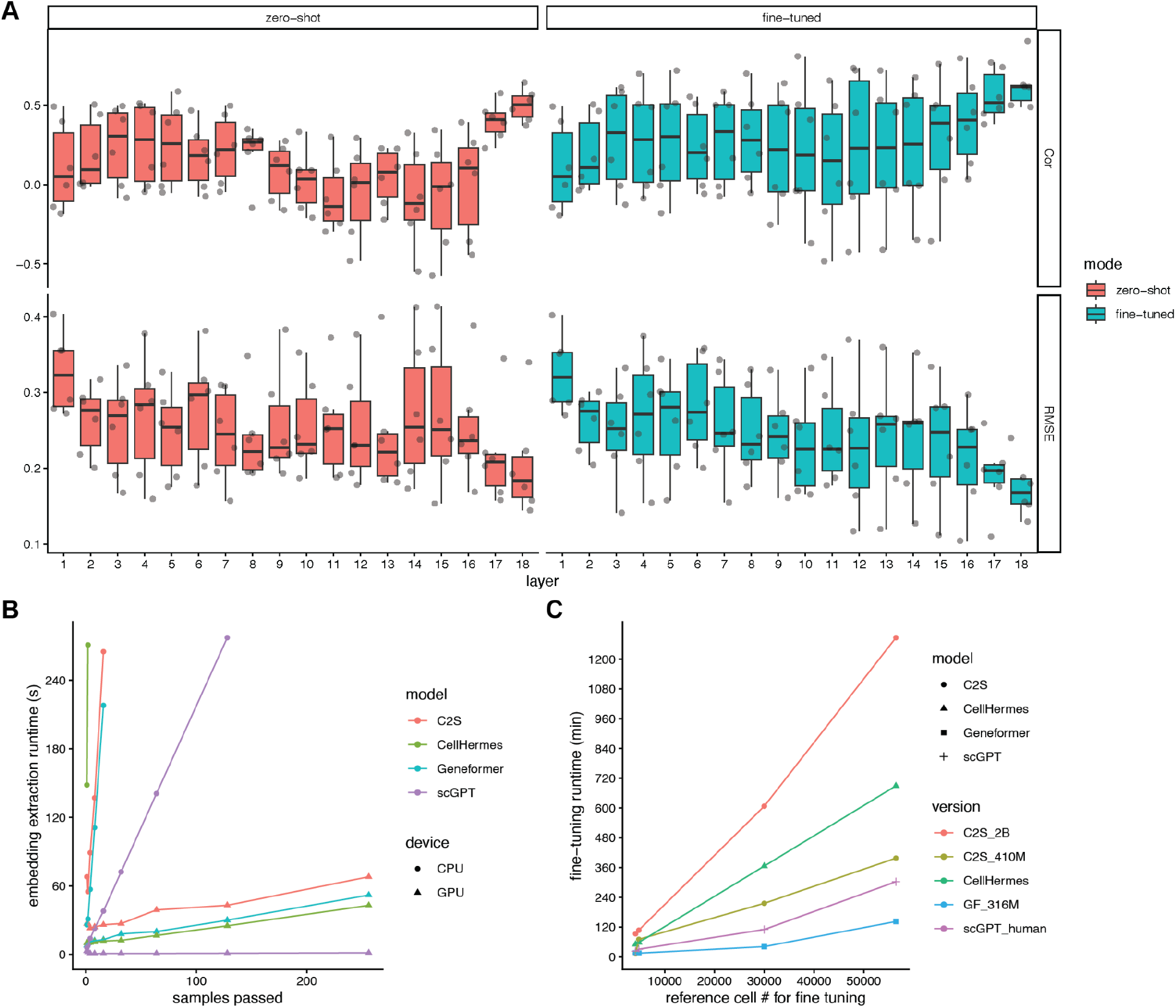
Technical details of DECONVersation implementation. A) Geneformer embedding extraction of the final layer showed better deconvolution performance than prior layers, using real bulk data with ground truths. B) Number of samples or cells scaling runtime comparison of various scFMs for embedding extraction on CPU or GPU. C) Cell number scaling runtime comparison of various scFMs for fine-tuning on GPU. **ALT TEXT**: Graphs labelled A to C detailing DECONVersation implementation performance. A is a box plot comparing deconvolution accuracy across Geneformer layers. B is a line graph showing CPU and GPU runtime scaling by sample or cell count. C is a line graph showing GPU runtime scaling during fine-tuning across cell numbers.

Despite the many utilities of scFMs, their evaluation and adoption have been limited by resource requirements, particularly GPU access. However, because the DECONVersation workflow reduces large single-cell references to signature matrices of a handful of cell types, and a deconvolution run often involves only tens to hundreds of RNA-seq samples, it is far less demanding than analyses of thousands to millions of single cells. We assessed the computational cost of generating scFM embeddings (**Fig 4B**). With GPU access, extracting embeddings from the signature matrix and bulk samples was fast, with all methods taking under 30 seconds for 10 samples. On CPU, most scFMs took under 270 seconds for 10 samples. In both cases, runtime scaled roughly linearly with the number of samples for all models, making projections straightforward. Fine-tuning increased total runtime, but all scFM methods tested took under 240 minutes for 10,000 cells (**Fig 4C**), with runtime scaling approximately linearly with the number of cells. Importantly, since the fine-tuning task is standard cell type classification, the fine-tuned models can also be used for other single-cell applications, such as integration or in silico perturbation. Likewise, if a model has been previously fine-tuned on a specific single-cell dataset for these applications, DECONVersation can reuse it directly for bulk deconvolution.

## DISCUSSION

Here, we extend single-cell foundation models to bulk RNA-seq deconvolution, asking whether pre-trained and fine-tuned embeddings can remain robust to technical variation, capture relevant biological signals, and preserve the linear additive relationships between cell types. We evaluated this across six real bulk datasets with ground-truth cell proportions from previous deconvolution benchmarks, spanning several tissues, and simulated pseudobulks that test finer cell type resolution and modality mismatch with the reference. Overall, scFM embeddings are surprisingly effective, given that these are general purpose models, trained on and for single-cell rather than bulk data. Although we focus on human scFMs, where the largest training corpora have been compiled, the same principles should apply to other organisms paired with corresponding foundation models.

scFM-derived embeddings generally outperformed raw gene expression for deconvolution. At broad cell type or lineage level, fine-tuned C2S-2B and Geneformer were consistently competitive with DWLS, the strongest conventional tool in our and recently published benchmarks, and outperformed other tested standard tools. For finer subtypes, we recommend fine-tuning, consistent with gains reported for cell-type labeling^42^. Yet the competitive performance of zero-shot Geneformer shows that cell type proportions can be recovered from the general embedding space without fine-tuning. Given the added cost of fine-tuning, zero-shot Geneformer offers the best accuracy-runtime trade-off for lineage or broad type-level deconvolution among the scFMs tested. Future developments in the scFM field, such as larger single-cell atlases and newer architectures, may further improve zero-shot deconvolution and simplify the deconvolution workflow.

Performance differences across scFMs suggest their embeddings encode different information.For example, scGPT and CellHermes performed worse on bulk PBMCs after fine-tuning. One explanation is that models differ in how well they separate cell identity from other variations in their training data, such as batch, donor, and assay. Gene rank ordering (Geneformer) and sentence ordering (C2S) may act as normalization that suppresses technical variation, explaining their stronger performance. Alternatively, because we reproduced each model’s recommended cell type classification fine-tuning, the fine-tuning processes are very different between models. For example, scGPT fine-tuning uniformly reduced accuracy slightly, possibly reflecting previously reported catastrophic forgetting^43^.

On pseudobulks drawn from the same single-cell source as the reference, every method (scFM-based, standard tools, and raw counts) exceeded 0.8 global correlation, suggesting any of these tools is appropriate when technical differences are minimal. The advantage of scFMs emerges as source-reference differences increase. Across the graded endometrial comparisons (same lab to different lab to different assay), correlation declined for nearly all methods, but less for scFM-based methods than for raw counts and standard tools. DWLS was the notable exception among conventional tools, holding performance across conditions.

As more scFMs emerge, tasks and metrics are needed to benchmark them against each other, raw expression, and conventional approaches. While previous benchmarks have focused on single-cell use cases such as perturbation and clustering, deconvolution may reveal how embeddings capture biological signal versus noise, cell type specificity, and quantitative representation of full transcriptomes. Advances in expression representation (e.g. TranscriptFormer’s^44^ expression-aware attention), architecture, and training corpora (larger collections or tissue-specific curations) may further improve deconvolution accuracy. A new class of foundation models trained on bulk RNA-seq data, such as BulkFormer^18^, have been reported to outperform scFMs on several bulk transcriptomic tasks, although deconvolution has not been evaluated. Single-cell reference-based deconvolution sits at a unique junction where the foundation model needs to represent both real bulk and single-cell derived cell-type pseudobulk signatures, yet neither scFMs nor bulkFMs were designed to do both. As DECONVersation is model agnostic, incorporating and evaluating BulkFormer is a natural next step.

Embedding generation for a signature matrix and bulk samples is efficient on a GPU, requiring under 100 seconds for 100 samples. While CPU extraction is substantially slower, we implement and test CPU-supported functions so DECONVersation on ∼10–30 samples completes in under an hour without a GPU. We also adapted Geneformer and C2S (410M and 2B) embedding extraction for Apple Silicon MPS acceleration, which took half or less the CPU time on a MacBook Air M2 (16GB). While CellHermes supports MPS natively, with similar speed improvements over CPU, its 8B-parameter V1 model exceeds 16GB RAM and is partially offloaded to disk, slowing extraction speed. Fine-tuning requires a GPU and considerable time, but scales approximately linearly with cell number and is comparable to intensive steps in other workflows such as DWLS marker selection or BayesPrism modeling on many bulk samples.

Moreover, a fine-tuned scFM supports many other analyses, including data integration, in silico perturbation, and network construction.

### Limitations

Our conclusions have several limitations. First, ground truths in the real bulk datasets were estimated by cell count mixtures, flow cytometry, imaging, or single-cell profiling, and their errors may extend to our results. Second, we cannot determine whether reference datasets overlap with scFM pretraining corpora, which are large and not always fully enumerated. Since the tested tissues (except tumor cell lines) are frequently profiled by single-cell technologies, larger models have likely seen relevant biology. As such, scFM zero-shot performance on entirely new tissues and rare cell types cannot be predicted. Third, all scRNA-seq references are annotation-dependent, and bias or errors in annotation (especially at the finer cell type labels) may affect all performance reported. For pseudobulk tests, we intentionally selected datasets integrated and annotated together, for label consistency. However, this too could be susceptible to other biases. Fourth, deep learning embeddings are hard to interpret or trace to specific genes. Although we provide tools to explore attention, attention weights alone do not always provide causal explanation^45^. As scFMs differ in training corpus, expression representation, and potential linearity distortion, it is difficult to isolate the factors driving deconvolution performance. Notably, even for standard deconvolution methods, benchmarking studies have struggled to explain why certain tools fare poorly on specific cell types or tissues^7,27^. Finally, although embedding generation is mostly parameter-free, it is limited by the default number of tokenized genes, effectively a top-expressed-gene filter on bulk profiles. While these defaults worked across our tissues and cell types, they may not suit other tissues in question.

## Supplemental Figures

**S1.**
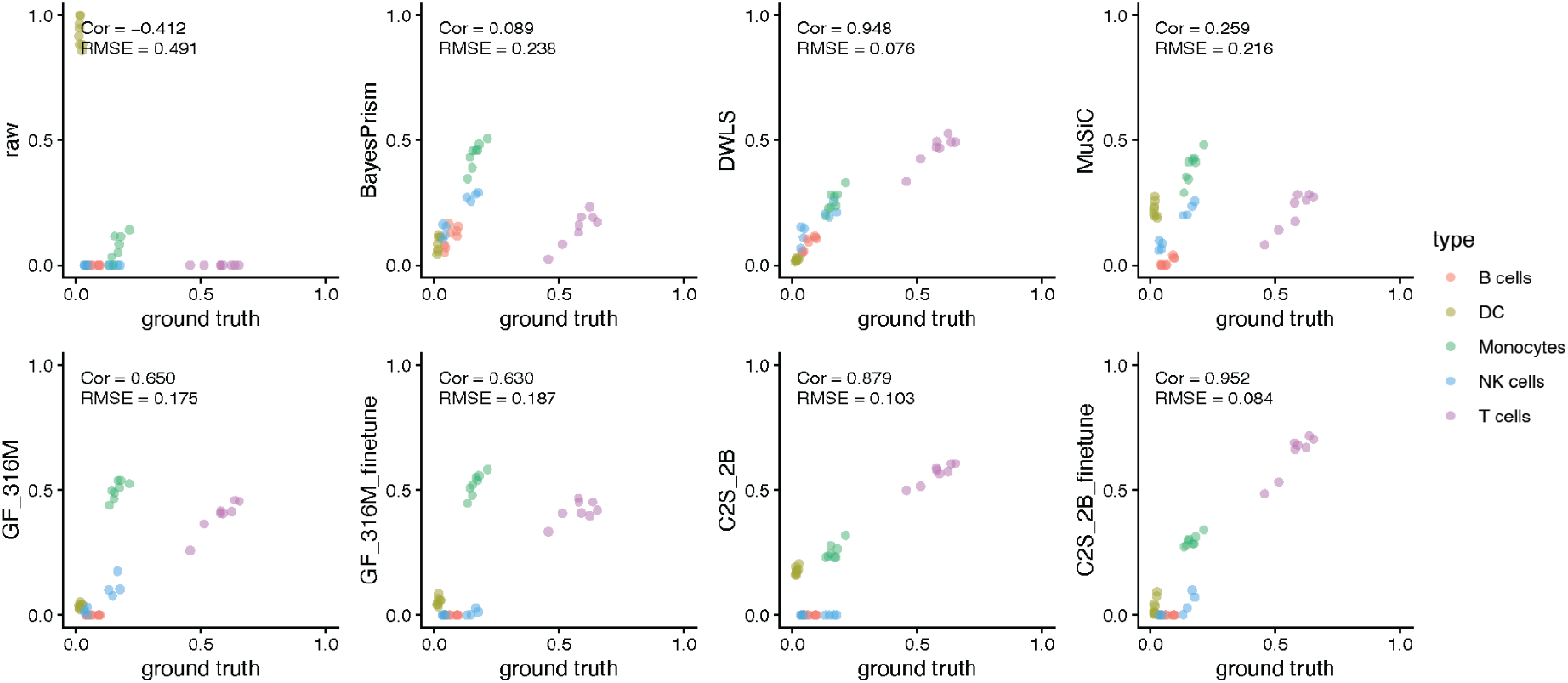
Example results from Hoek et al. PBMC bulk dataset, with cell sorter counts as ground truth vs predictions from different tools.

**S2.**
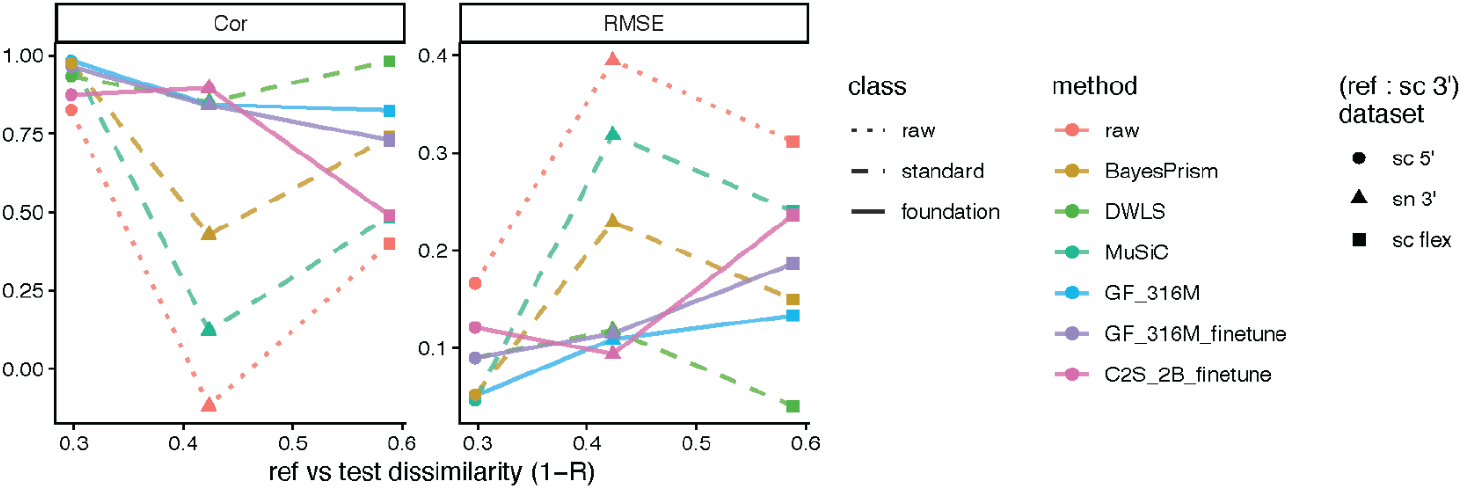
Comparison of various deconvolution tools using the same PBMC single cell 3’ reference against increasingly dissimilar single cell/nuclei datasets.

**S3.**
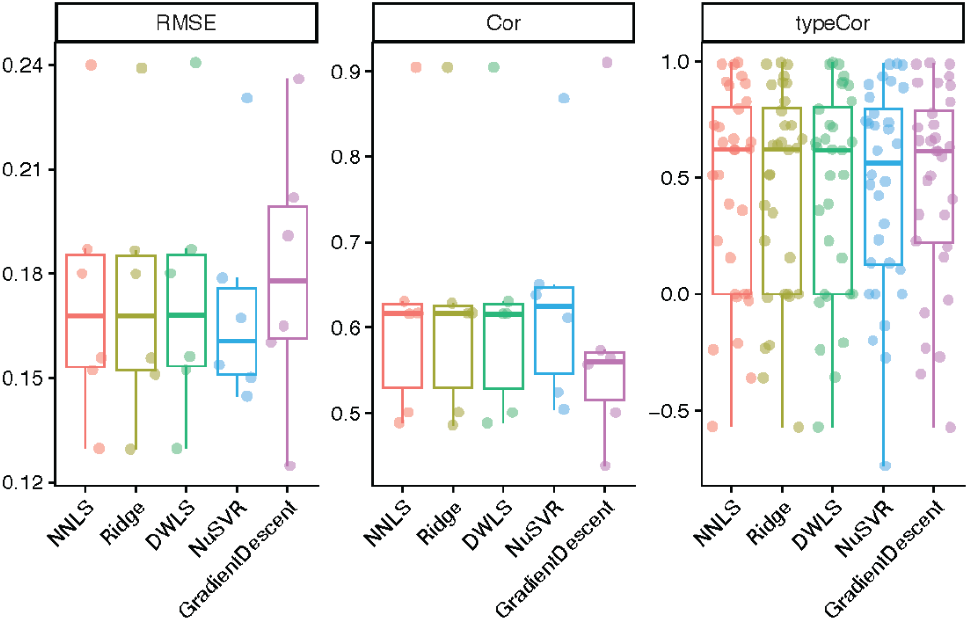
Comparison of NNLS and other solvers on Geneformer zero-shot embeddings on real bulk data.

## Code availability

DECONVersation code and tutorials are available at https://github.com/nygenome/DECONVersation and can be installed from PyPI at https://pypi.org/project/DECONVersation/.

## Data availability

Bulk and pseudobulk datasets, corresponding ground truths, and single-cell reference objects are archived at https://zenodo.org/records/22949405

## Acknowledgements

We thank Dr Will Liao and Dr Jiaxi Liang for discussions.

We used large language models, ChatGPT and Claude, to assist with code and package unit test generation. All outputs were critically reviewed and edited by the authors, who take full responsibility for the final content.

