## Supplemental Figs 1,2,3 for "DECONVersation: Single-Cell Foundation Model-Derived Embeddings for Robust Cell Type Deconvolution of Bulk RNA-seq"

Supplemental Figures

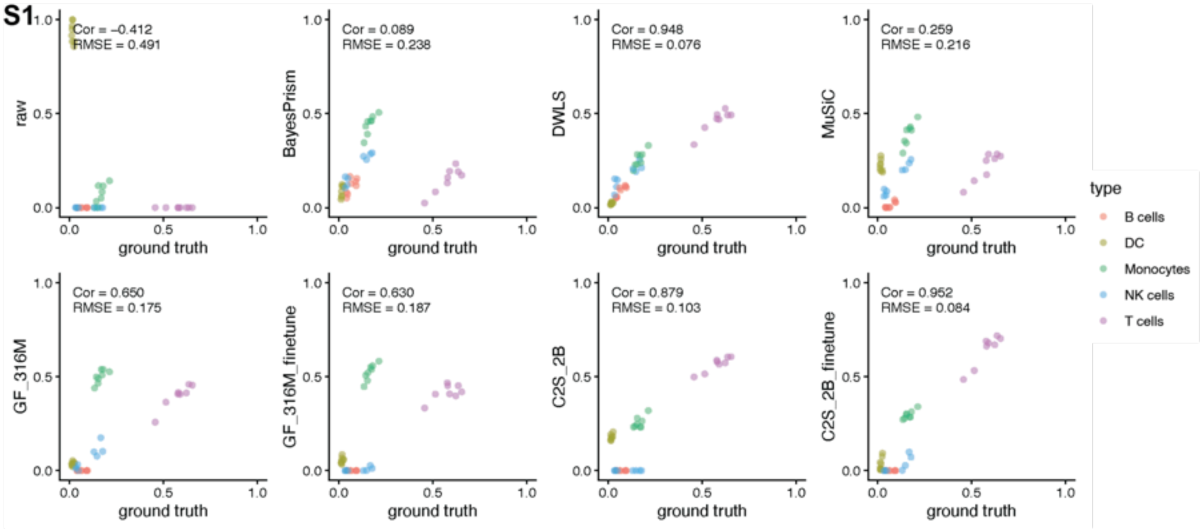


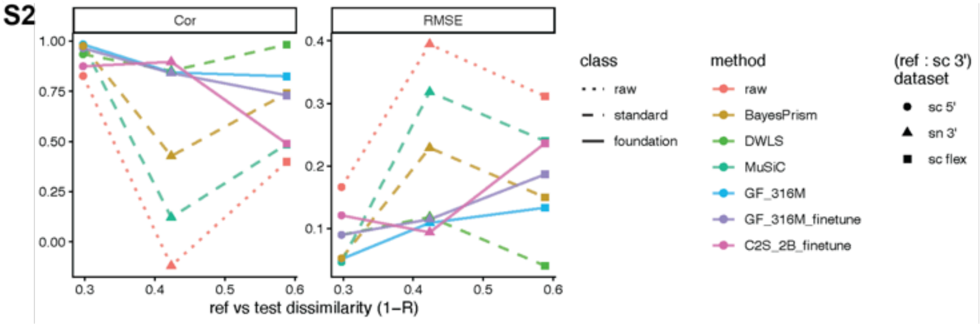


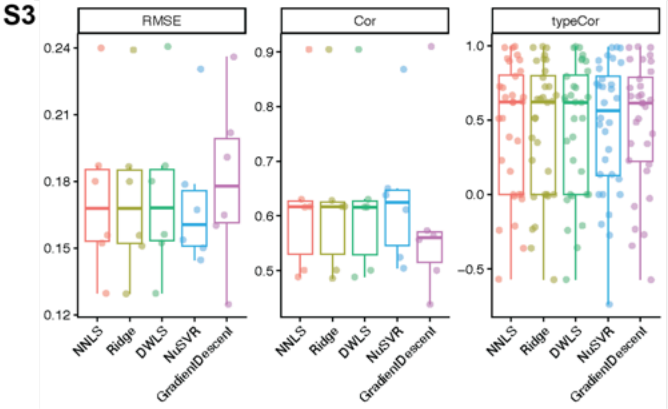


S1. Example results from Hoek et al. PBMC bulk dataset, with cell sorter counts as ground truth vs predictions from different tools.

S2. Comparison of various deconvolution tools using the same PBMC single cell 3' reference against increasingly dissimilar single cell/nuclei datasets.

S3. Comparison of NNLS and other solvers on Geneformer zero-shot embeddings on real bulk data.
